# ISCO: Intelligent Framework for Accurate Segmentation and Comparative Analysis of Organoids

**DOI:** 10.1101/2024.12.24.630244

**Authors:** Jian Zhou, Zihao Fu, Xiaoying Ni, Qi Luo, Gen Yang

## Abstract

Organoids are self-organizing 3D cell clusters that closely mimic the structure and function of in vivo tissues and organs. Quantifying organoid morphology is critical for advancing our understanding of organ development, drug discovery, and toxicity assessment. Recent advances in microscopy have provided powerful tools to capture detailed morphological features of organoids, yet manual image analysis remains labor-intensive and time-consuming. In response, we present a comprehensive microscopy-based analysis pipeline that utilizes SegmentAnything 2.1 to accurately segment individual organoids. Additionally, we introduce a suite of morphological features—including perimeter, area, radius, non-smoothness, and non-circularity—that enable researchers to quantitatively and automatically analyze organoid structures. To further standardize organoid analysis across the field, Intelligent segmentation and comparison of organoids(ISCO), an intelligent AI-driven open-source algorithm, is developed with the aim of establishing a comprehensive toolset for organoid characterization.

## Introduction

Organoids are self-organizing, three-dimensional cell aggregates derived from stem cells through in vitro differentiation, capable of recapitulating the structure and function of native tissues. As an innovative model for cancer research, organoids faithfully replicate the physiological characteristics of in situ tissues, preserving the essential features of tumor cells in vivo. Their ability to mimic complex biological processes positions organoids as a versatile platform with broad applications in biomedical research, including developmental and disease modeling, drug toxicity testing, gene and cell therapy evaluation, and drug screening. These properties render organoids a promising tool for advancing the field of biomedical science and technology.

Recent advancements in organoid research have been significantly propelled by the development of deep learning platforms, such as OrganoID, which automate the identification, annotation, and tracking of individual organoids in microscopy experiments. This platform demonstrates exceptional accuracy in monitoring organoid morphology and size — critical parameters for evaluating drug responses in high-throughput settings. Notably, OrganoID maintains tracking precision above 89% over extended periods, underscoring its potential to accelerate organoid-based research and enhance its application in drug discovery and other biomedical investigations. Concurrently, significant progress has been made in the development of high-throughput image analysis tools designed to handle the complexity and volume of data generated by organoid experiments. One such tool, Cellos, offers an advanced image analysis pipeline for 3D organoid and nuclear segmentation. By combining classical algorithms with a Stardist-3D convolutional neural network, Cellos enables accurate segmentation of organoids and nuclei, facilitating detailed analysis of cellular dynamics and drug responses, particularly in cancer pharmacology. Similarly, OrBITS integrates computer vision and machine learning to provide automated live-cell imaging, enabling kinetic monitoring of organoids and enhancing drug screening processes.

The integration of machine learning into organoid analysis has greatly improved both the functionality and accessibility of these tools. For example, MOrgAna, a Python-based software, leverages machine learning for image segmentation and morphological analysis of organoids. It is designed to be user-friendly, requiring no coding expertise, while also offering customization options for advanced users, making it suitable for a wide range of biomedical applications. Similarly, OrganoID utilizes deep learning to track and analyze single-organoid dynamics, providing accurate and detailed image analysis in high-throughput contexts.Additionally, specialized software such as Organalysis has been developed for specific organoid types, such as cardiac organoids, offering multifaceted image preprocessing and analysis capabilities, including noise removal, feature importance computation, and fractal analysis. These tools enhance the precision and versatility of organoid research, particularly in regenerative medicine. Moreover, a novel large model-derived algorithm for complex organoids, focused on internal morphogenesis and digital marker derivation, outperforms existing methods in evaluating intricate organoid structures.In summary, the latest advancements in organoid analysis emphasize high-throughput capabilities, machine learning integration, and user-friendly interfaces, which have collectively enhanced the efficiency and accuracy of organoid data analysis. Tools such as Cellos, MOrgAna, OrBITS, and OrganoID represent key milestones in the field, providing robust solutions for diverse organoid applications and driving forward research in drug discovery and disease modeling.

### Workflow

Organoid images were acquired using an inverted microscope with a 10X objective lens. For each sample, 5-8 random fields were captured at 1024×980 pixel resolution and saved in png format. Image preprocessing included Gaussian filtering (σ=1.5) and contrast enhancement using CLAHE algorithm. Organoid segmentation was performed using a U-Net-based deep learning model, followed by watershed algorithm for separating connected organoids. The model was trained on manually annotated organoid images and achieved a mean Intersection over Union (IoU) of 0.91.Quantitative analysis of organoid morphology was conducted using custom Python scripts. Key measurements included area (μm^2^), perimeter (μm), circularity, and aspect ratio. The segmentation results were validated through manual inspection of randomly selected samples. Statistical analysis was performed in R (version 4.1.0), using Shapiro-Wilk test for normality assessment and one-way ANOVA for group comparisons. P-values less than 0.05 were considered statistically significant.

The analysis pipeline generated comprehensive reports containing growth curves, morphological parameter distributions, and statistical comparisons between experimental groups. Quality control measures included focus checking, illumination uniformity assessment, and batch effect correction. The entire workflow was implemented using Python 3.8 with OpenCV and PyTorch libraries for image processing and deep learning, while R was used for statistical analysis and visualization.

We defined the morphological parameters based on the SAM4organoid algorithm(Figure 2 and Table1 ).

**Figure 1.**
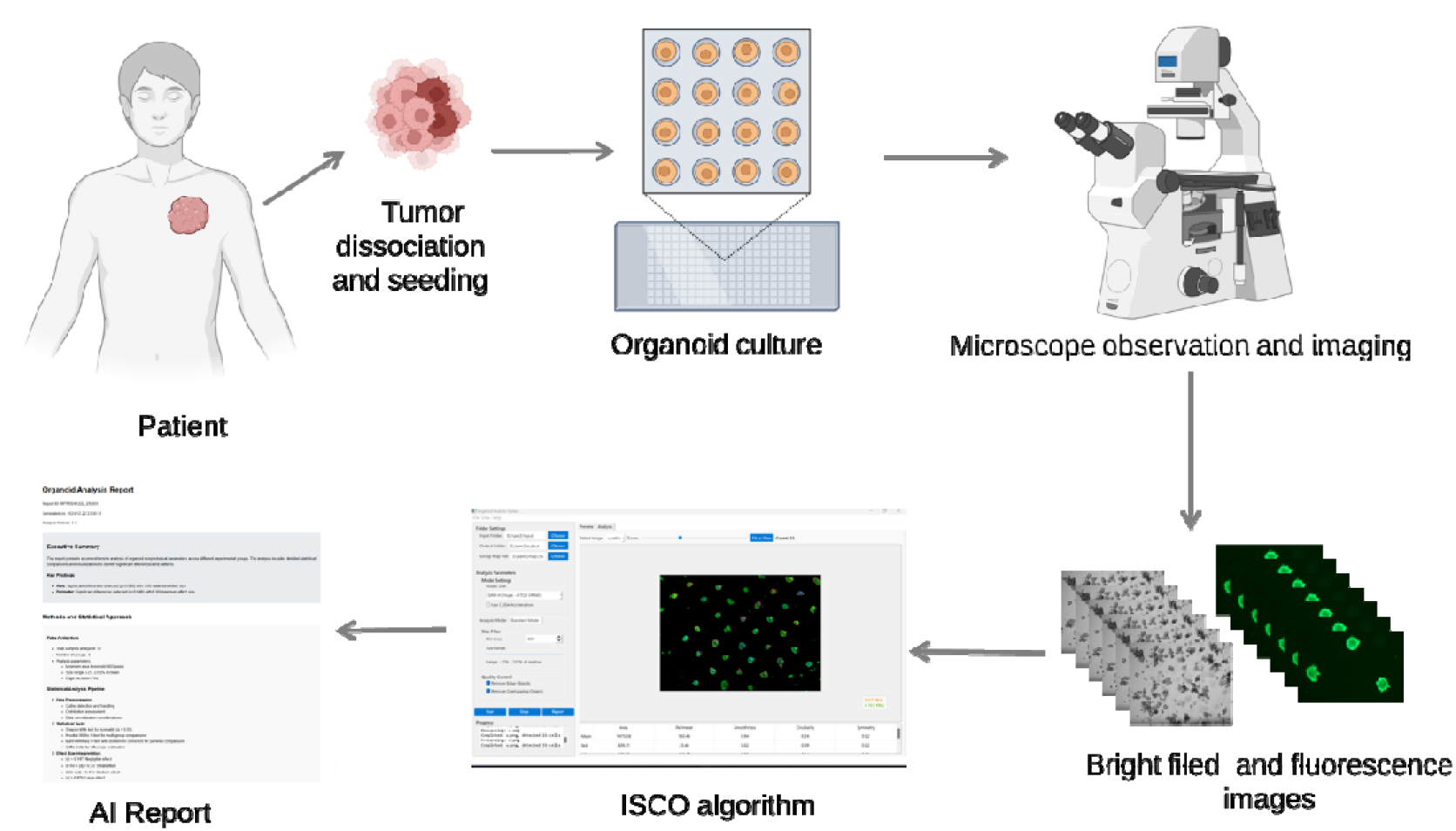
The process of tumor organoid culture establishment and intelligent reporting.

**Figure 2.**
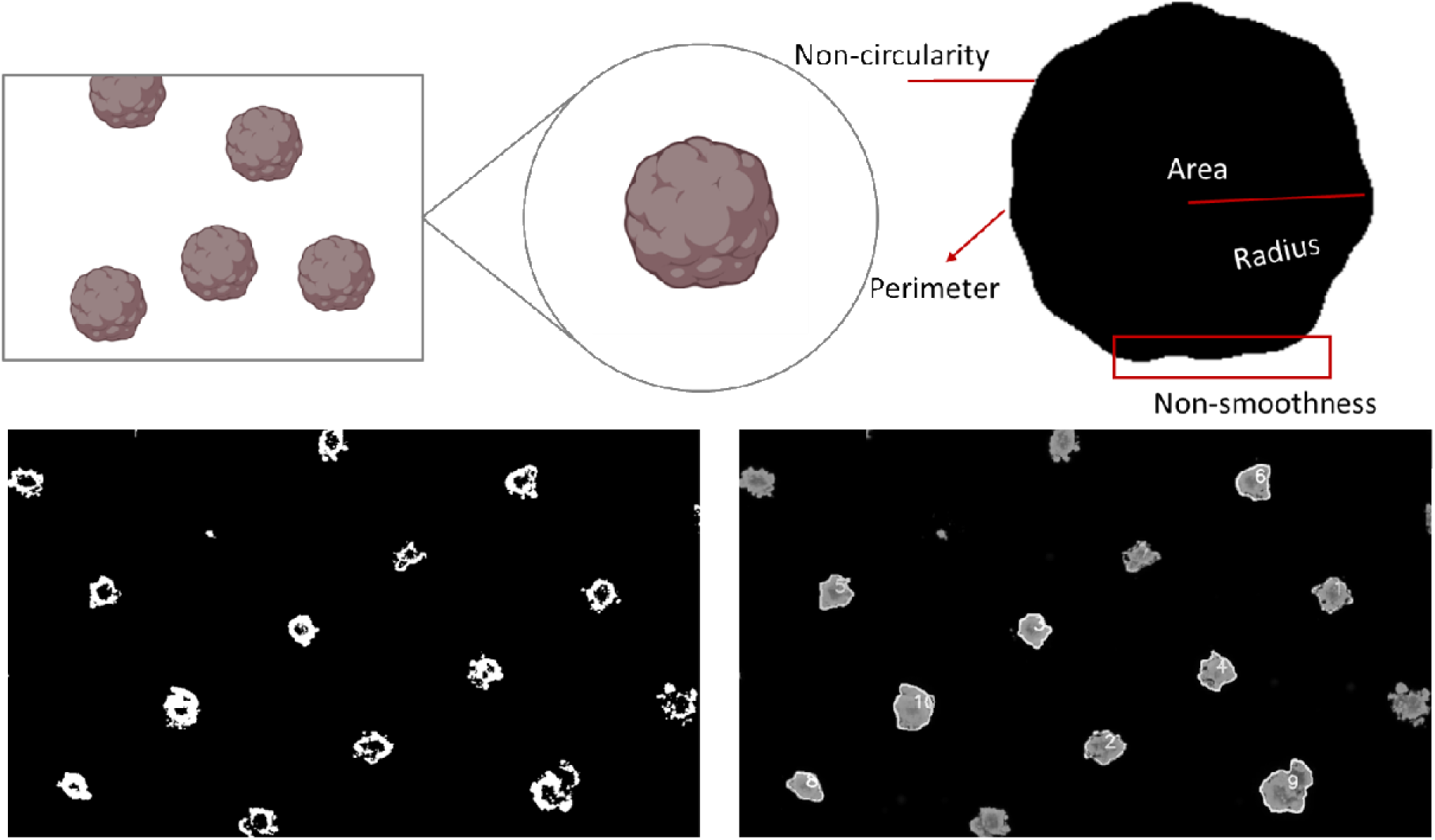
Organoid morphological parameters

**Table 1.**
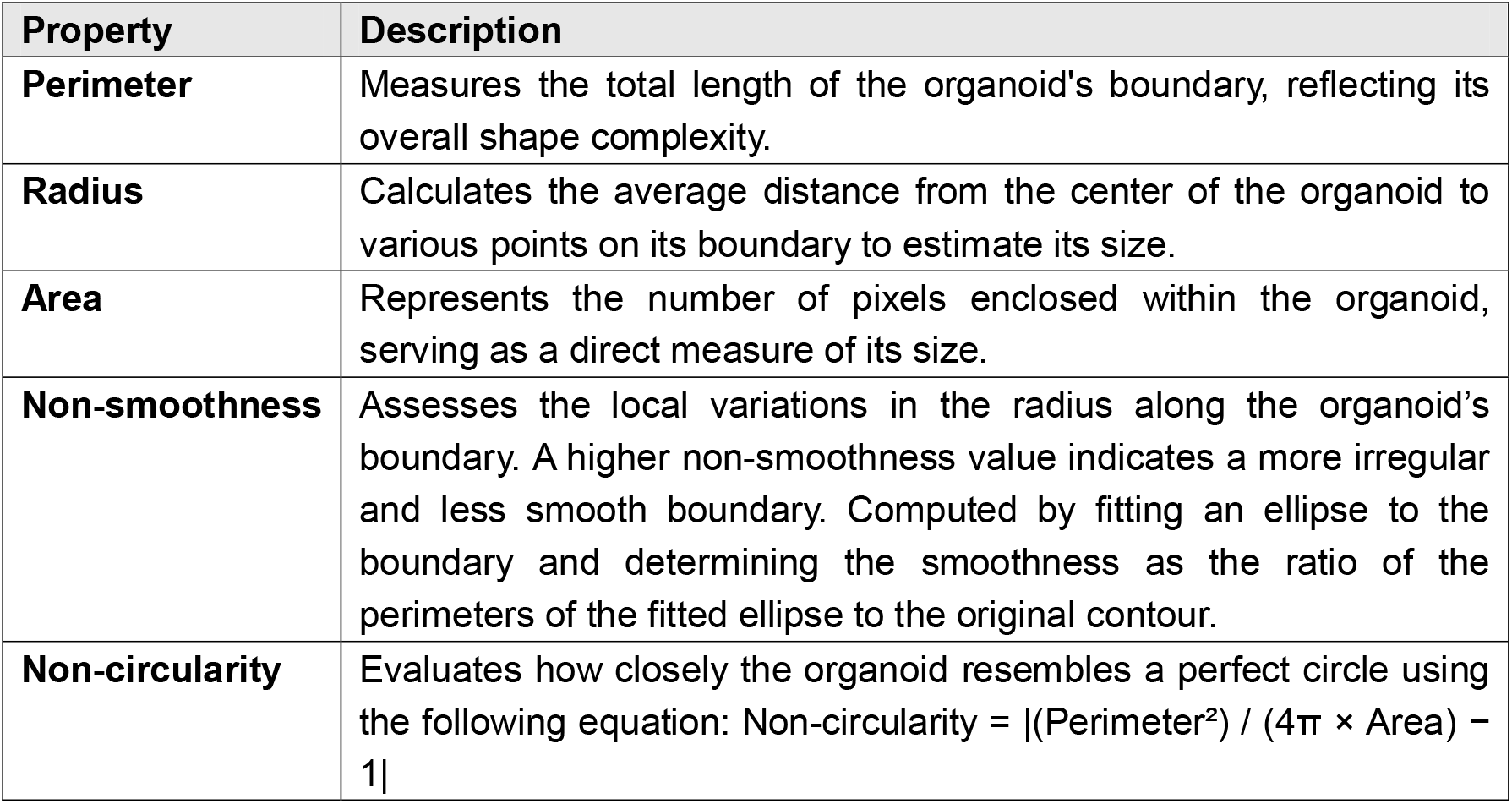
Organoid morphological parameters and their definitions.

We defined the morphological parameters based on the SAM4organoid algorithm (Figure 2 and Table 1). This algorithm provides a robust framework for quantifying key shape features of organoids, ensuring consistency and accuracy in parameter extraction. The parameters we calculated include perimeter, radius, area, non-smoothness, and non-circularity, which together offer a comprehensive description of the organoid morphology. These metrics enable a detailed comparison across different organoids, allowing us to assess variations in shape, size, and regularity.

As shown in Figure 2, the organoid boundaries were processed using the SAM4organoid algorithm, which incorporates advanced image processing techniques such as contour detection and ellipse fitting. Table 1 summarizes the specific formulas and methodologies used to compute each morphological property, with a particular focus on non-circularity, which was calculated using the equation:

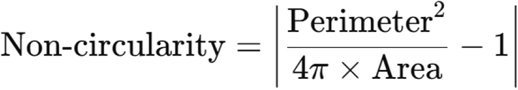

This approach ensures that all parameters are derived consistently and provides a clear framework for future comparisons. By applying this algorithm, we can achieve reliable and reproducible measurements of organoid morphology, facilitating further analysis in cellular behavior, growth patterns, and other biological studies.

Figure 3 shows the interface of the ISCO algorithm software, which was used to process and analyze the organoid images. The software provides a user-friendly platform for extracting morphological parameters such as perimeter, radius, area, non-smoothness, and non-circularity, as defined by the SAM4organoid algorithm. The interface allows for easy input of organoid images, which are automatically processed to detect boundaries, calculate the required metrics, and visualize the results.

**Figure 3.**
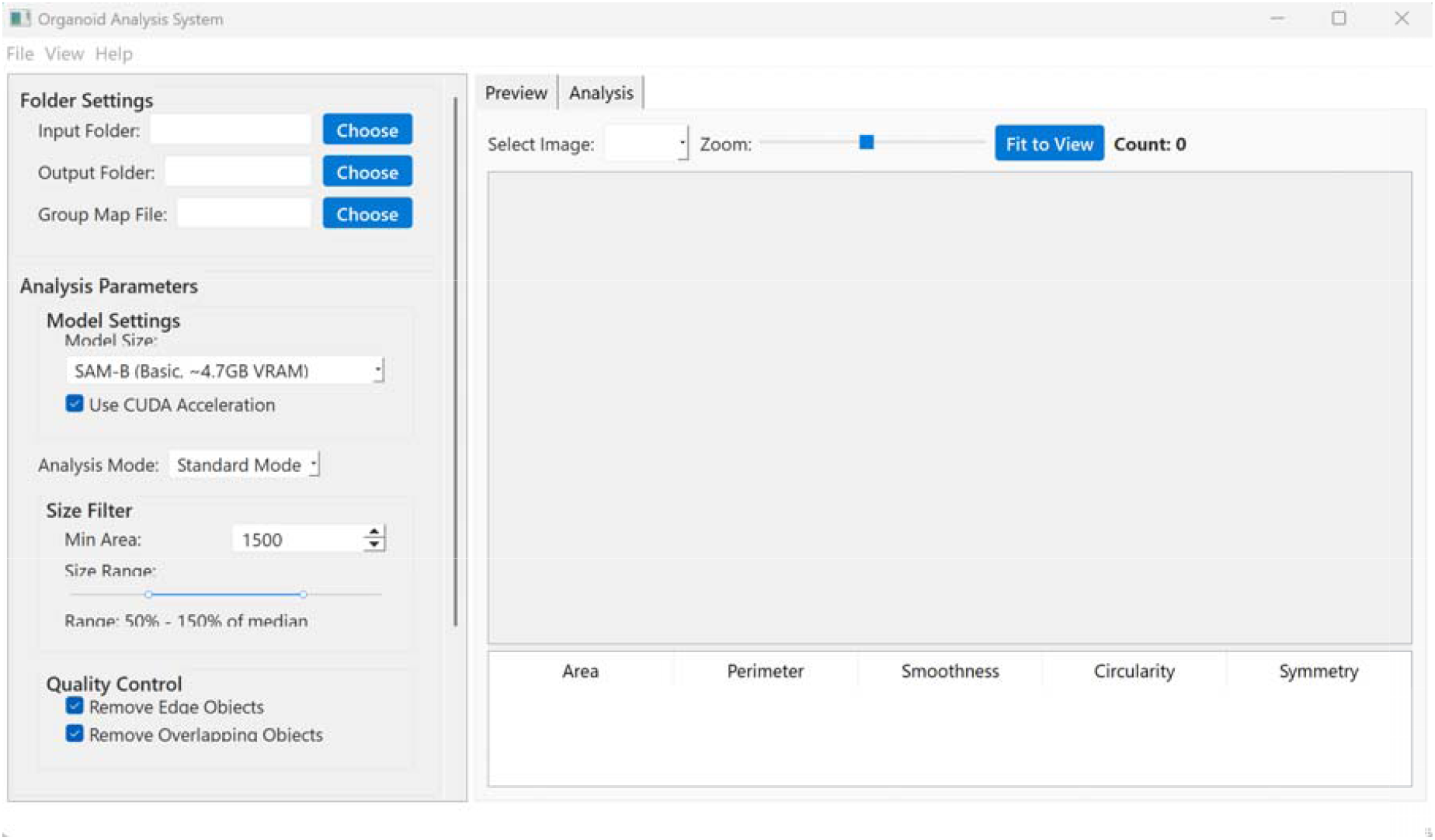
the interface of the ISCO algorithm software

Using ISCO, we were able to efficiently analyze large sets of organoid images, minimizing human error and ensuring the reproducibility of measurements. The software’s output includes both numerical data and graphical representations, such as the contour overlays and scatter plots, which allow for a visual comparison of organoid shape and size across different samples.The integration of the ISCO algorithm with SAM4organoid’s morphological parameter definitions offers an enhanced approach for organoid characterization, enabling more precise and standardized analysis. This platform will also be useful for comparing the effects of different treatment conditions on organoid growth and morphology, providing valuable insights into cellular behavior and tissue development.

Figure 4 presents a violin plot case from the group comparison analysis generated by the ISCO algorithm. This plot provides a visual representation of the distribution of morphological parameters, allowing for easy comparison of the variability and central tendency of the data across different groups. By utilizing violin plots, we can better understand the range of values for parameters such as perimeter, radius, area, non-smoothness, and non-circularity, and identify any significant differences between experimental groups or treatment conditions.The ISCO algorithm’s ability to produce such comparative visualizations not only enhances data interpretation but also facilitates statistical analysis, enabling researchers to identify subtle trends and outliers. The integration of these visualization tools into the algorithm significantly improves the ease of analyzing complex datasets, supporting more informed decision-making in experimental design and interpretation.

**Figure 4.**
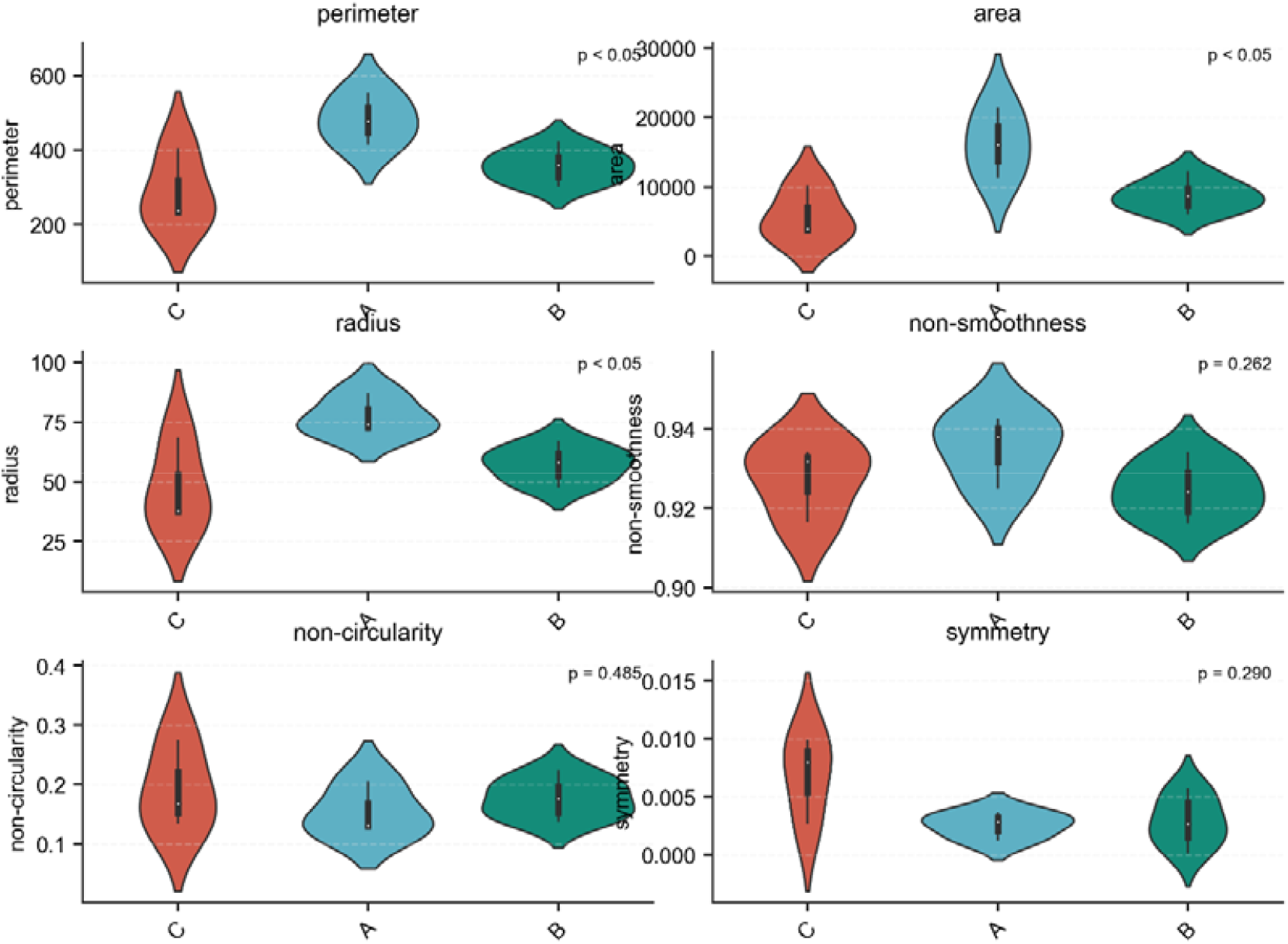
a violin plot case from the group comparison analysis generated by the ISCO algorithm

### Future Directions

Looking ahead, the ISCO algorithm holds significant potential for further development. One possible direction is the integration of machine learning techniques, such as deep learning-based segmentation, to improve the accuracy of boundary detection, especially in challenging or low-resolution images. Additionally, incorporating more advanced statistical tools for multi-dimensional data analysis could further enhance its capabilities in assessing organoid morphology and other biological characteristics.

The open-source nature of the ISCO algorithm plays a crucial role in its potential for continuous improvement and widespread adoption. By making the algorithm freely available, the research community can contribute to its refinement, identify and fix bugs, and implement new features that address specific research needs. Open-source development fosters collaboration, enables transparency, and accelerates the adoption of advanced computational tools across different scientific disciplines. Furthermore, it empowers researchers with the flexibility to adapt and customize the algorithm to suit their specific applications, whether in organoid research, tissue engineering, or other fields involving morphological analysis.

